# First evidence of *in vitro* cytotoxic effects of marine microlitter on *Merluccius merluccius* and *Mullus barbatus*, two Mediterranean commercial fish species

**DOI:** 10.1101/2021.11.10.468019

**Authors:** A Miccoli, E Mancini, PR Saraceni, G Della Ventura, G Scapigliati, S Picchietti

**Affiliations:** Department for Innovation in Biological, Agro-food and Forest Systems, University of Tuscia, Viterbo, 01100, Italy; Italian Fishery Research and Studies Center, Rome, 00184, Italy; Department of Science, Roma 3 University, Rome, 00146, Italy; INFN Laboratori Nazionali di Frascati, Via E. Fermi 54, Frascati, 00044, Italy

**Keywords:** Marine microlitter, Bioindicators, Cytotoxicity, *In vitro* approaches, Primary cell cultures, Biological agents

## Abstract

Marine litter is composed mainly of plastics and is recognized as a serious threats to marine ecosystems. Ecotoxicological approaches have started elucidating the potential severity of microplastics (MPs) in controlled laboratory studies with pristine materials but no information exist on marine environmental microlitter as a whole. Here, we characterized the litter in the coastal Northern Tyrrhenian sea and in the stomach of two fish species of socio-economic importance, and exposed primary cell cultures of mucosal and lymphoid organs to marine microlitter for evaluating possible cytotoxic effects. An average of 0.30 ± 0.02 microlitter items m^-3^ was found in water samples. μFT-IR analysis revealed that plastic particles, namely HDPE, polyamide and polypropylene were present in 100% and 83.3% of *Merluccius merluccius* and *Mullus barbatus* analyzed, which overall ingested 14.67 ± 4.10 and 5.50 ± 1.97 items/individual, respectively. Moreover, microlitter was confirmed as a vector of microorganisms. Lastly, the apical end-point of viability was found to be significantly reduced in splenic cells exposed *in vitro* to two microlitter conditions. Considering the role of the spleen in the mounting of adaptive immune responses, our results warrant more in-depth investigations for clarifying the actual susceptibility of these two species to anthropogenic microlitter.

**Highlights:**

- 0.30 ± 0.02 microlitter items m^-3^ were found at the surface of coastal Northern Tyrrhenian sea
- 14.67 ± 4.10 and 5.50 ± 1.97 items/individual were retrieved from the stomach of hakes and mullets
- The ingested microlitter contained plastic items
- Microlitter was validated as a carrier of bacteria, fungi and flagellates
- Splenic cells exposed to two microlitter conditions for 72 hours suffered cytotoxicity

## 1. Introduction

Coastal areas are subject to an exponential increase in population density and the development of impacting human activities, e.g. industries, tourism, recreational activities, fishing and aquaculture. As a consequence, they can be affected by both sporadic and continuous pollution events, with consequences on all compartments, and are thus considered “hotspots” of contamination (Llorca et al., 2020; Lu et al., 2018).

Marine litter is defined as “any anthropogenic manufactured, or processed solid material (regardless of size) discarded, disposed of, or abandoned in the environment, including all materials discarded into the sea, on the shore, or brought indirectly to the sea by rivers, sewage, storm water, waves, or winds” (UNEP, 2016). It is one of the most serious threats to marine ecosystems and a global environmental concern. It comprises glass, metal, cardboard and textiles items (Löhr et al., 2017) as well as anthropogenic particles produced by industrial activities (e.g., coal-fired power plants) and transport emissions (Piazzolla et al., 2020), but Tekman et al. (2021) revealed that plastic accounts for the 66-79% of the global litter composition.

Annual global plastic production accounted for 368 million tons in 2019 (Association of Plastic Manufacturers, 2020): about 1.3-3.1% of these (5-12 million tons year^-1^) reach the Oceans (Jambeck et al., 2015), and the total amount of floating plastic was estimated at 0.3 million tons (van Sebille et al., 2015). The Mediterranean Sea, in particular, is regarded as an accumulation zone for marine litter, with densities comparable to those of the five subtropical gyres (Cózar et al., 2015; UNEP/MAP, 2015; Van Sebille et al., 2020).

In addition to primary microplastics (MPs), i.e. particles that are purposefully manufactured of microscopic sizes < 5 mm, the vast majority of marine litter is subject to degradation by abiotic (UV radiation, mechanical abrasion, temperature) and biotic (microbiological depolymerization) agents, resulting into secondary MPs (Ru et al., 2020; Thompson et al., 2004). Their chemico-physical properties, such as type of polymer, density, size, shape, internal geometry and color, influence their transport, buoyancy and sinking as well as rates of ingestion and removal by aquatic organisms (Kowalski et al., 2016; Nguyen et al., 2020; Shim et al., 2018).

Due to their small size, MPs are bioavailable to a variety of taxa (e.g. Cole et al., 2013; Fossi et al., 2018; Gomiero et al., 2018; Lusher et al., 2013; Pittura et al., 2018) and can either be mistaken with or selectively chosen instead of food (Clark et al., 2016; Moore, 2008), with demonstrated impacts. Once ingested, MPs can affect biological functions and tissue integrity of organisms (Cole et al., 2015; Pedà et al., 2016; Sussarellu et al., 2016). Moreover, MPs can be potential carriers of pollutants (Amelia et al., 2021; Guo and Wang, 2019) and can be colonized by microbial pathogens, transferring them along the trophic web (Caruso, 2019; Casabianca et al., 2019). Ecotoxicological and physiological impacts of MPs were also demonstrated in controlled laboratory studies, but the commercially-available and pristine materials employed hardly reflect the actual heterogeneity of the environmental litter.

The European hake *Merluccius merluccius* (Linnaeus, 1758) and the red mullet *Mullus barbatus* (Linnaeus, 1758) are good experimental models in MPs-related research because of their biological features, commercial relevance, abundance in the Mediterranean region and suitability as small-scale plastic pollution bioindicators. Building on research that demonstrated the presence of MPs in their gastrointestinal tracts (Atamanalp et al., 2021; Avio et al., 2020, 2015; Bellas et al., 2016; Digka et al., 2018; Giani et al., 2019; Mancuso et al., 2019), in the present study we hypothesized that microlitter could be retrieved from the marine environment and ingested by fish, and that it could affect the *in vitro* viability of immunologically-relevant cells. We therefore investigated the litter abundance in the coastal sector of Civitavecchia (Northern Tyrrhenian Sea, Latium, Italy) at the sea surface and in the stomachs of *M. merluccius* and *M. barbatus*. We also evaluated the microbiological contamination and the *in vitro* cytotoxicity of environmentally-collected microlitter particles on primary cell cultures of mucosal and lymphoid organs, with the ultimate aim of defining the actual susceptibility of these two socio-economically important fish species to anthropogenic microlitter.

## 2. Material & Methods

### a. Study area

The research was conducted in the northern Tyrrhenian Sea (Italy), FAO’s General Fisheries Commission for the Mediterranean (GFCM) Geographical Sub-Area 9 (GSA 9). The experimental campaign fell within the physiographical unit (PU) M. Argentario – Cape Linaro, and included the coastal platform that extends from Santa Severa (42.01676 N, 11.95604 E) to the Tarquinia coastal area (42.22243 N, 11.70495 E). The study area is characterized by a 120-150 m deep continental margin and a sandy to sandy-muddy seabed, as reported in Mancini et al. (2021).

### b. Experimental campaign

The experimental campaign took place on October 23^rd^ 2020 and extended over a 14-hour period (03:00 - 17:00 UTC/GMT +2:00). Four 15-minute long horizontal tows with a 250 µm mesh size net were performed at sea surface for litter sampling (Fig. 1). Table 1 reports time of start, GPS position, micro- and macrolitter abundance m^-3^ and meteo-marine conditions as per the World Meteorological Organization sea state coding.

**Fig. 1.**
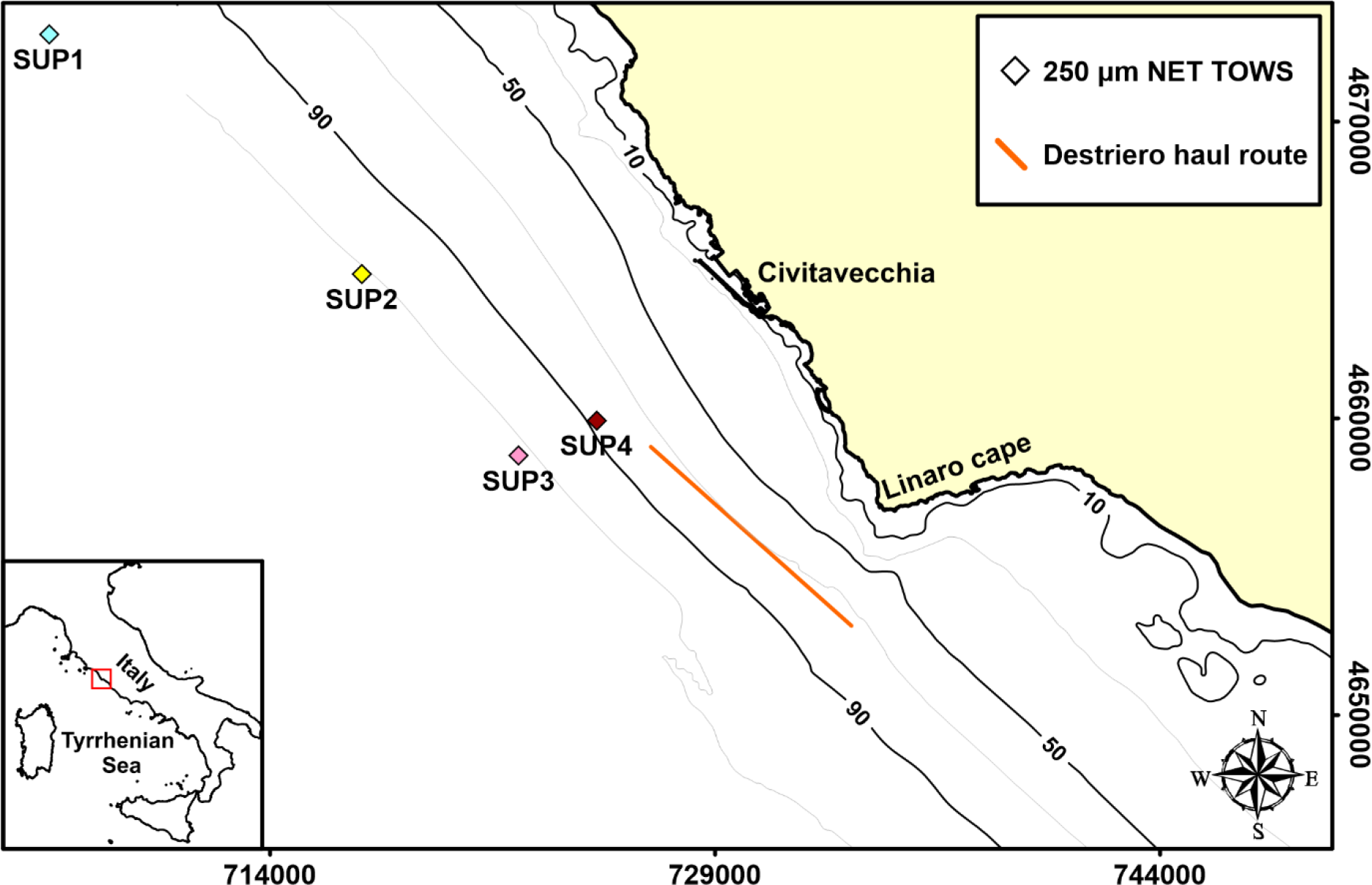
Map of sampling sites - Study area with 250 µm net tows GPS positions, haul route and bathymetry.

**Table 1.**
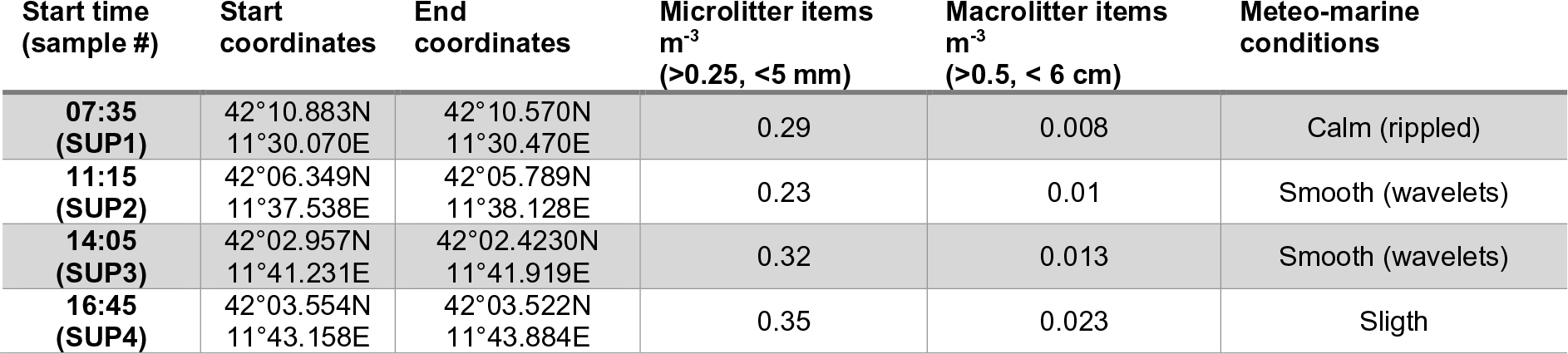
Sampling details - Summary of the experimental campaign with net tows details. Time expressed as UTC/GMT +2:00.

### c. Qualitative and quantitative characterization of litter at the sea surface

The 250 µm WP2 net was equipped with a metered line, a non-filtering cod-end and a flow meter. Upon deployment from the starboard side of the vessel, utmost care was taken to ensure that the net stretched out correctly. Upon retrieval, the content of the cod-end was transferred to 500 mL containers and stored at 4 °C until transported to the laboratory. To prevent airborne contamination in the laboratory, microlitter exposure to air was kept as short as possible and all following steps were performed under a sterile laminar flow cabinet previously cleaned with 100% ethanol. Each sample (SUP1-SUP4) was visually sorted 5 mL at a time using a stereomicroscope (Leica 8APO). Microlitter was classified in terms of shape (i.e. filament, fragment, film), color and diameter size (A: 250 < x < 500 μm; B: 500 < x < 1000 μm; C: 1000 < x < 3000 μm; D: 3000 < x < 5000 μm; E: x > 5000 μm). Size fractionation, as a method for representing litter size, was used in other recent peer-reviewed publications (Digka et al., 2018; Giani et al., 2019). Following the processing in the lab, microlitter sampled from the marine environment was stored in sterile 50 mL Falcon tubes filled with 0.22 µm-filtered sea water. Microlitter abundances are presented as items m^-3^ of seawater ± standard error (SE).

### d. Fish sampling and characterization of litter quantity and quality in stomach contents

Fish specimens were collected in the same area (October 30^th^ 2020, 01:11:00 PM haul start time, haul start GPS position 41°59.669 N 11°49.214 E, haul end GPS position 42°03.0410 N 11°44.4621 E) by means of the bottom trawl net typically known as “volantina” geared with a cod-end mesh size of 50 mm diamond and a vertical opening of 4 m (Sala et al., 2013). *M. merluccius* and *M. barbatus* specimens had a total length (TL) of 18.6 ± 1.02 cm and 11.7 ± 0.64 and a body weight (BW) of 47.1 ± 8.7 g and 16.5 ± 1.15 g, respectively (average ± SD): they were employed for the quali-quantitative characterization of microlitter within the stomach contents (n=6 per species) and for cytotoxicity assays (n=3 per species). Immediately following the opening of the net, specimens of similar within-species TL were sorted, thoroughly rinsed with 0.22 µm-filtered Milli-Q water, wrapped in autoclaved aluminum foil and immediately transferred to ice, where they were kept until lab processing. Stomach eversion was not observed during sampling.

In the laboratory, fish stomachs were excised under a sterile laminar flow cabinet, previously cleaned with 100% ethanol, and preserved in 75% ethanol until processing. Tools for organ dissection (e.g. stainless steel tweezers and forceps, glass Petri dishes) were sterilized by autoclave. Stomach contents were then placed in sterile Petri dishes and visually sorted with a stereomicroscope (Leica 8APO) to classify litter particles in terms of shape, diameter size and color, using the same classification as in section 2c. The particles retrieved from the stomach contents were maintained in 50 mL falcon tubes with 0.22 µm-filtered sea water. Microlitter abundances are presented as mean particles per species ± standard error (items ± SE). Our workflow was compliant with the recommendations of Bessa et al. (2019) with regards to species selection, sampling methods, airborne/external cross-contamination prevention and particle classification.

### e. μFT-IR analysis

To ascertain the nature of the microlitter items retrieved from fish stomach contents, forty items were analyzed by Fourier-transform infrared microspectroscopy (μFT-IR) following visual inspection. The subsample was representative of shape types, color and size ranges of items both within- and between-species. The experiments were performed at the DAFNE Laboratory of INFN (Frascati, Italy) in transmission mode, using a Bruker Hyperion 3000 FTIR microscope equipped with a Globar IR source, a broadband beamsplitter (KBr) and a mercury-cadmium telluride (MCT) detector; the beam size was set at 20×20 μm. Spectroscopic analysis yielded absorbance spectra, which were analyzed using the Open Specy open source database (Cowger et al., 2021) with the Pearson’s correlation coefficient as measure of the linear correlation between the data sets. Spectra visualization and overlay were achieved with the SpectraGryph 1.2 software using the peak normalization method (i.e. each spectrum highest peak within the visible area was set to 1).

### f. Assessment of microlitter cytotoxic effects on fish primary cell cultures

Marine microlitter particles were obtained by surface 250 µm net tows from the same sampling location of fish, and were stored in 50 mL falcon tubes filled with 0.22 µm-filtered sea water following visual inspection until employed for cytotoxicity assays. Microlitter particles were dried under laminar flow cabinet on a lint-free tissue (Kimwipes, Kimtech Science, USA) in a sterile glass Petri dish at room temperature and there counted using a stereo microscope (size ranging from 250 and 5000 µm, colors observed red, blue, black, green, grey). Isolation and cultivation of fish primary cells were performed according to published standard procedure (e.g. Miccoli et al., 2021b). Gills (G), head kidney (HK) and spleen (SPL) were dissected from *M. merluccius* and *M. barbatus* (n=3) under a sterile laminar flow cabinet previously cleaned with 100% ethanol with sterilized tools (see section 2d), and immediately immersed in cold Hank’s Balanced Salt Solution without calcium and magnesium (HBSS), previously adjusted for appropriate sea water osmolarity (355 mOsm Kg^−1^) with 3M NaCl. Cells were obtained through teasing in cold HBSS using disposable strainers of 100 µm and 40 µm mesh size, and homogenates were washed by centrifugation (10 min, 400 g, 4 °C). Cells were then resuspended in L-15 (Leibovitz) medium containing 10% heat-inactivated fetal calf serum (FCS, Gibco) and antibiotics (penicillin–streptomycin, Gibco). Cells were counted in a Neubauer chamber and adjusted to a concentration of 5×10^5^ cells mL^-1^ in L-15 medium. Both HBSS and L-15 media were 0.22 µm-filtered.

Cellular suspensions were exposed to “Low” and “High” microlitter conditions corresponding to 4 and 20 field-collected microlitter particles/ml. Both microlitter concentrations were consistent number-wise with the ingestion rates of microlitter herein described, and the lowest microlitter concentration was in line with future modelled estimates weight-wise (Isobe et al., 2019).

Cells were cultured for 2 and 72 hours at 15 °C with gentle rotary shaking to ensure a continuous contact with microlitter particles. Because the bottom trawl net was not equipped with any temperature sensor, the incubation temperature was selected according to the near real-time numerical model MEDSEA_ANALYSISFORECAST_PHY_006_013 (Clementi et al., 2021), resolving for variable “sea_water_potential_temperature_at_sea_floor (bottomT)” using sampling location, date and depth as input. Three technical replicates per biological sample were used in all experimental groups. Negative controls consisting of cells incubated at the same conditions without microlitter were considered for each organ and species. Positive controls, i.e. cells incubated as negative controls but with 0.2% NaN3, were also tested separately for each organ and species. Intracellular ATP value, as a proxy of cell viability/cytotoxicity (Schoonen et al., 2005), was then quantitatively evaluated simultaneously in all samples of both species for any given incubation time using the ATPlite assay (PerkinElmer, catalog no. 6016943) following the manufacturer’s instructions: 50 µL of cell lysis and 50 µL substrate solutions were added to 100 µL cell suspensions per replicate and shaken for 5 min. Resulting homogenates were transferred to disposable opaque well plates (OptiPlate-96, PerkinElmer) and luminescence was measured using a microplate reader (Wallac Victor2, PerkinElmer), following a 10-minute dark adaptation period.

### g. Validation of microlitter as a carrier of biological agents

At the end of a 72-hour incubation of primary cultures with microlitter following an alike experimental design as above, 10 μL of cell suspension from each experimental group were qualitatively observed under a Zeiss microscope equipped with a colour 8 camera (AxioCam MRC) and a software package (KS 300 and AxioVision). Multiple sets of photographs at random frames were taken per each experimental group and microorganisms were quantified over a 100.000 µm^2^ area by an operator unaware of treatments. Sterile glass slides and cover slips were used for these steps.

### h. Data analysis, visualization and statistics

Stomach content particle abundance was tested for statistical significance between species using an independent samples t-test with the null hypothesis of equal population means between groups. Datasets were checked for normality with the Shapiro-Wilk test and for homoscedasticity with the Levene test. A log-transformation was applied to meet the normal distribution assumption.

The relation among species and ingestion of microlitter particle types by color was examined with a chi-square test on a two-way contingency table. The null hypothesis assumed no association between variables. Results are reported as χ^2^_df_ =test statistic.

Cytotoxicity data, grouped by species, time, organ and treatment, were tested for statistical significance using a one-way ANOVA with the null hypothesis of equal population means among groups, followed by a Tukey’s HSD *post-hoc* test in case the main effect of the models was significant. Datasets were checked for normality with the Shapiro-Wilk test and for homoscedasticity with the Levene test.

Microbial count was analyzed with the rank-based nonparametric Kruskal-Wallis test because datasets, many of which were zero-inflated, did not meet the assumptions for parametric testing. The null hypothesis was that samples were drawn from the same population or from populations whose medians did not differ.

A comparison of microlitter densities retrieved from representative scientific literature was visualized as mean ± SD items m^-3^ with the R “forestplot” package v 2.0.1. Data reported in other units than items m^-3^ were excluded from the analysis. References were organized hierarchically by location and year, and box size was set to constant.

### i. Ethics statement

Animal manipulation complied with the guidelines of the European Union Directive (2010/63/EU) and the Italian Legislative Decree 26 of 4 March 2014 “Attuazione della Direttiva 2010/63/UE sulla protezione degli animali utilizzati a fini scientifici”. Ethical review and approval was not required because animals were sampled from the natural environment and were not subject to any experimental manipulation, in line with the Explanatory Note of the Italian Ministry of Health’s Directorate-General for Animal Health and Veterinary Medicinal Products (DGSAF) of 26 July 2017.

## 3. Results

### a. Qualitative and quantitative characterization of litter at the sea surface

Anthropogenic marine litter visually of plastic origin and in the form of filaments, fragments and films was found in all water samples taken in the Civitavecchia area (Fig. 2, Table 1). Both microlitter (250 μm < x < 5 mm) and macrolitter (5 mm < x < 6 cm) fractions were identified. SUP4 and SUP2 had the highest and lowest microlitter particle densities with 0.35 and 0.23 items m^-3^, respectively (Table 1). SUP4 and SUP 1 had the highest and lowest macrolitter particle densities with 0.023 and 0.008 items m^-3^, respectively (Table 1). Their average abundance among all samples was 0.30 ± 0.02 items m^-3^ and 0.014 ± 0.003 items m^-3^ (Fig. 3A).

**Fig. 2.**
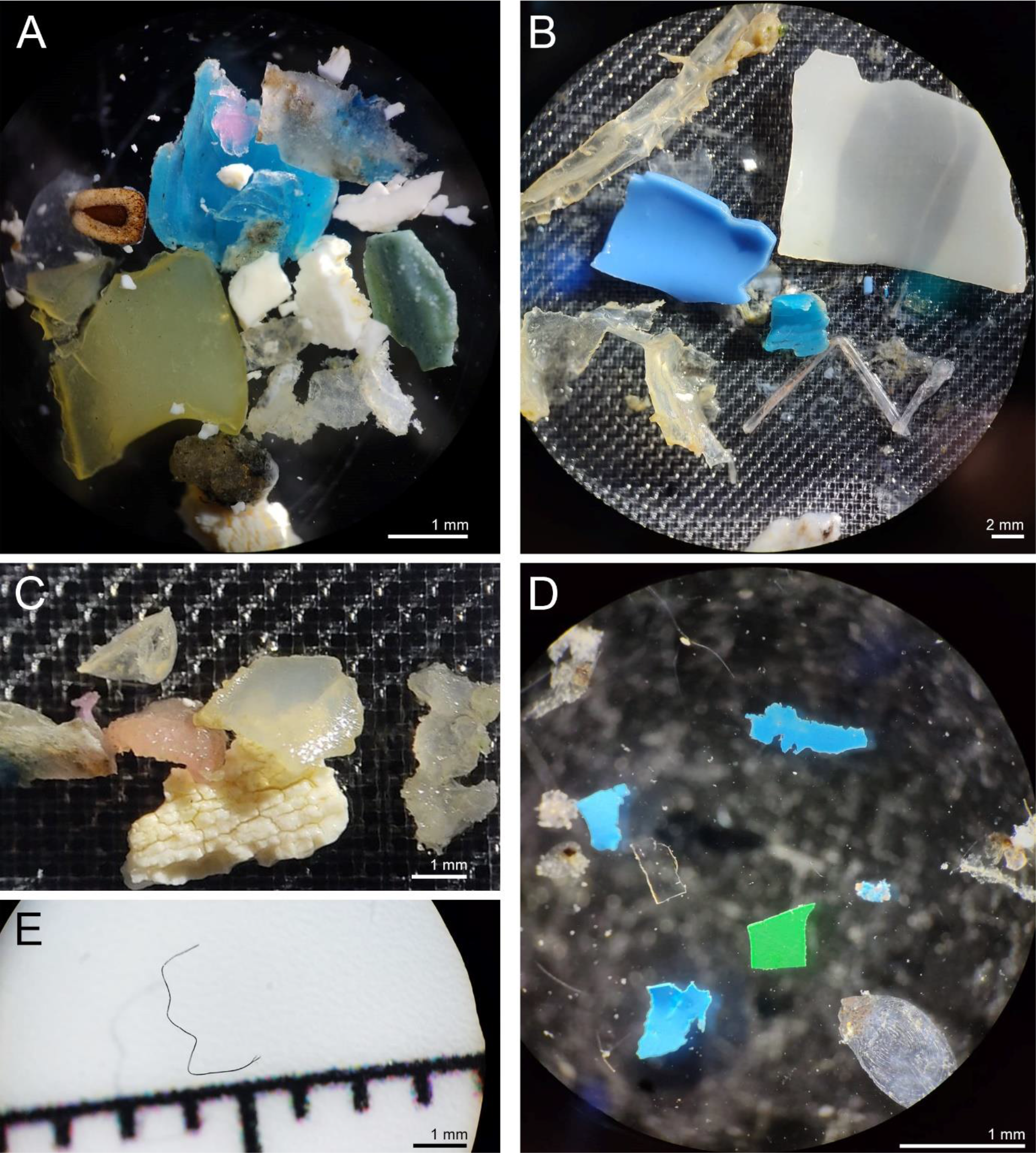
A-E Microlitter collected from sea surface - Some examples of litter particles collected from surface 250 μm net tows (SUP1-SUP4).

With regard to microlitter shapes, filaments, fragments and films were observed. In particular, the highest and lowest concentration of filaments were found in SUP4 (0.12 items m^-3^) and SUP3 (0.02 items m^-3^), respectively; the highest and lowest concentration of fragments were found in SUP3 (0.17 items m^-3^) and SUP1 (0.039 items m^-3^), and the highest and lowest concentration of films was found in SUP4 (0.18 items m^-3^) and SUP2 (0.07 items m^-3^) (Fig. 3B). Number and density of particles per type per color per station, total number of particles collected and relative share of litter types are reported in Table S1.

**Fig. 3.**
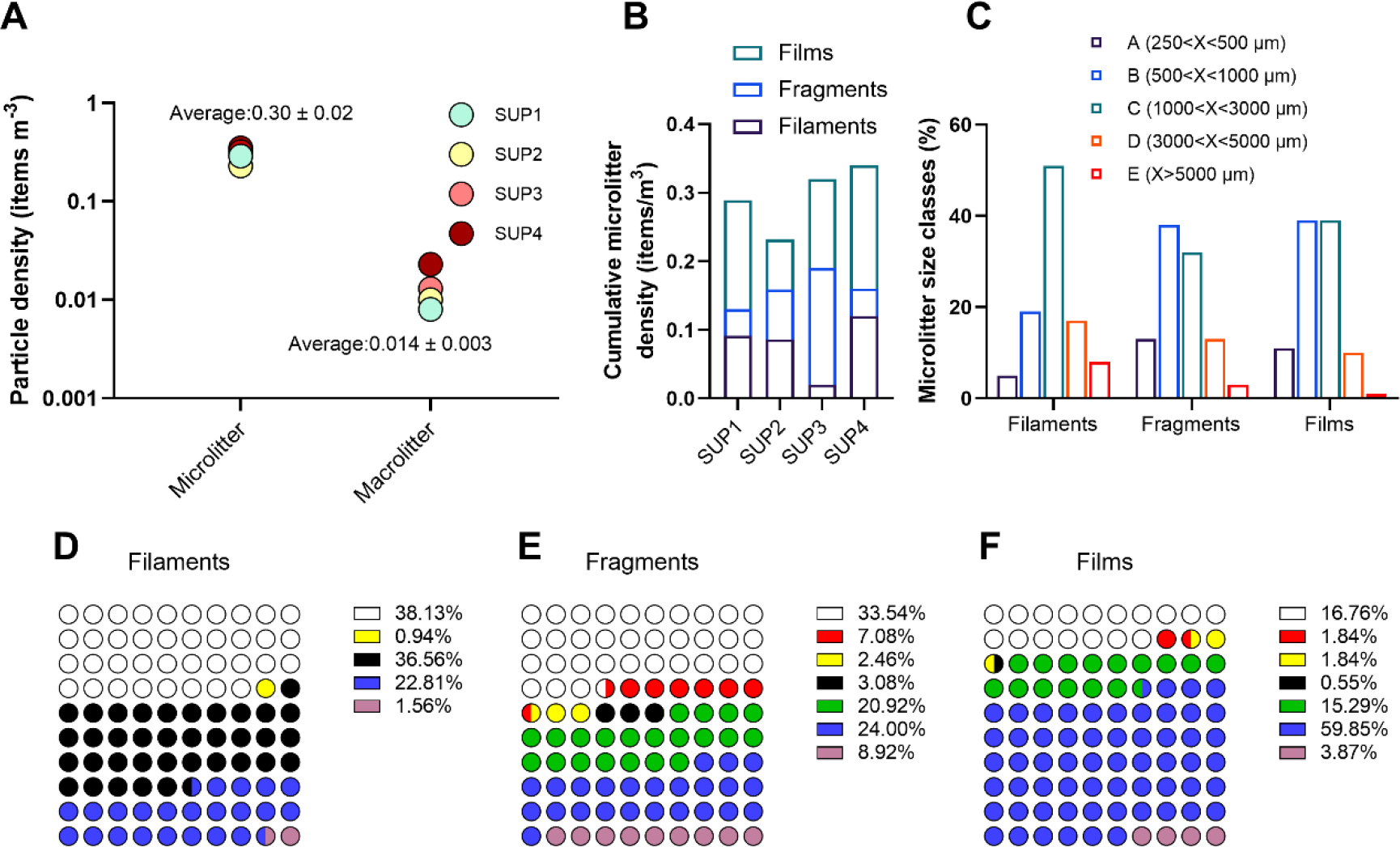
Quali-quantitative characterization of anthropogenic litter collected from the sea surface- A) Particle density of the micro- and macrolitter fractions per net tow. Average litter density is reported as mean ± SE. B) Cumulative microlitter density per item type per net tow. C) Size class distribution per litter shape type. D-F) Percentage of color abundance per microlitter shape type.

The vast majority (51%) of filaments belonged to size class C and a smaller population to size class A (5%). Fragments fell for the most part within size class B (38%), while size class E was the least represented (3%). Films mostly belonged to size classes B and C (39%), while size class E was found in the 1% of cases (Fig. 3C and Table S2).

**Table 2.** Full statistical details of the rank-based nonparametric Kruskal-Wallis tests performed on microbial counts per species and organ. ns: non significant.

| Species | Organ | Mean microbial count |  |  | SE microbial count |  |  | 95% CI microbial count [L;U] |  |  | Statistical test |  |  |
| --- | --- | --- | --- | --- | --- | --- | --- | --- | --- | --- | --- | --- | --- |
|  |  | ctrl | low | high | ctrl | low | high | ctrl | low | high | H | p |  |
| *M. merluccius* | Gills | 0 | 269 | 241.2 | 0 | 143.8 | 57.95 | 0;0 | -349.8;887.8 | -8.14;490.5 | 5.609 | 0.068 | ns |
|  | Head kidney | 0 | 14.4 | 23.31 | 0 | 7.00 | 3.91 | 0;0 | -15.79;44.48 | 6.49;40.13 | 6.161 | 0.025 | * |
|  | Spleen | 0 | 8.07 | 48.42 | 0 | 5.6 | 34.97 | 0;0 | -16.02;32.16 | -102.0;198.9 | 5.162 | 0.1 | ns |
| *M. barbatus* | Gills | 0 | 0 | 18.83 | 0 | 0 | 4.11 | 0;0 | 0;0 | 1.15;36.51 | 7.624 | 0.036 | * |
|  | Head kidney | 0 | 0 | 17.04 | 0 | 0 | 5.88 | 0;0 | 0;0 | -8.26;42.34 | 7.624 | 0.036 | * |
|  | Spleen | 0 | 5.38 | 23.31 | 0 | 5.38 | 13.92 | 0;0 | -17.77;28.53 | -36.58;83.21 | 4.587 | 0.107 | ns |

Filaments from all tows were mostly white (38%) and black (37%); less frequent colors in terms of abundance were blue (23%) and yellow (1%). Fragments were more chromatically diversified: while most of them were white (34%), blue and green fragments were found with a percentage of 24% and 21%, respectively; red (7%), black (3%) and yellow (2%) fragments were less abundant. Films were mostly blue (60%), followed by white (17%) and green (15%); yellow (2%) and red (2%) films were less frequent (Fig. 3D-F and Table S1).

### b. Characterization of quantity, quality and chemical composition of microlitter in fish stomach content

Microlitter items were found in 100% *M. merluccius* specimens and in 5 out of 6 (83.3%) *M. barbatus* specimens (Fig. 4).

**Fig. 4.**
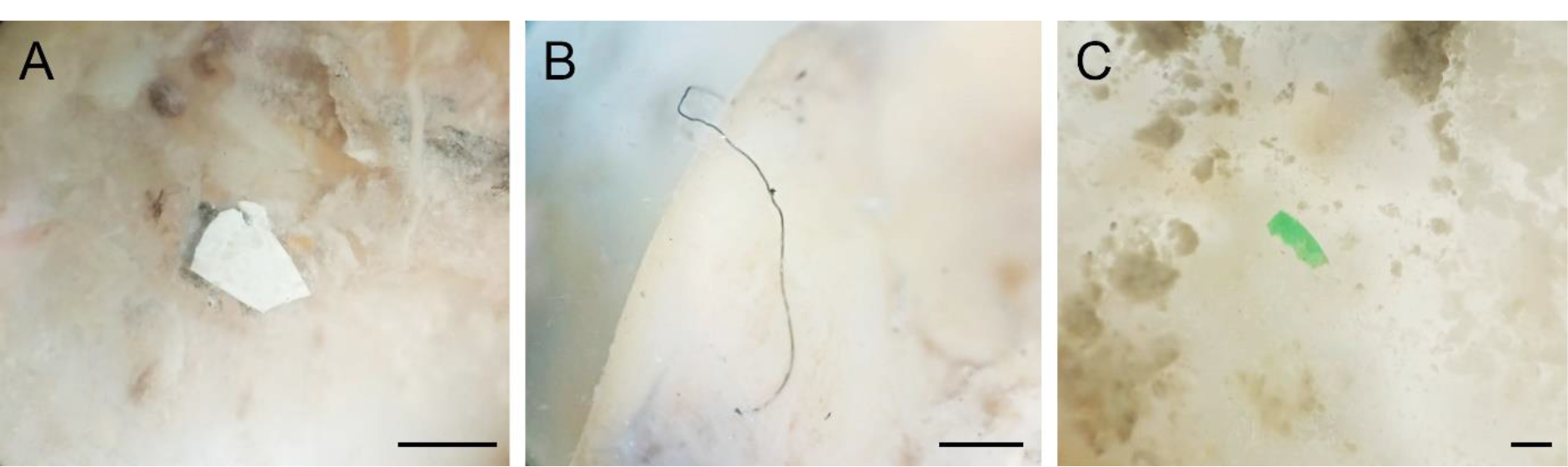
Examples of microlitter items retrieved from the digestive tract of fish - A) White fragment from *M. barbatus*, 0.58 match with polyamide (Primpke et al., 2018). B) Black filament from *M. merluccius*, 0.46 match with HDPE (Chabuka and Kalivas, 2020). C) Green fragment from *M. merluccius*, 0.72 match with HDPE (Chabuka and Kalivas, 2020). Scale bars: 500 μm.

A higher abundance of microlitter was found in the stomach contents of hakes (14.67 ± 4.10 items/individual) than mullets (5.50 ± 1.97 items/individual), but the difference between group means was not statistically significant (t10=2.045, p=0.068) (Fig. 5A).

**Fig. 5.**
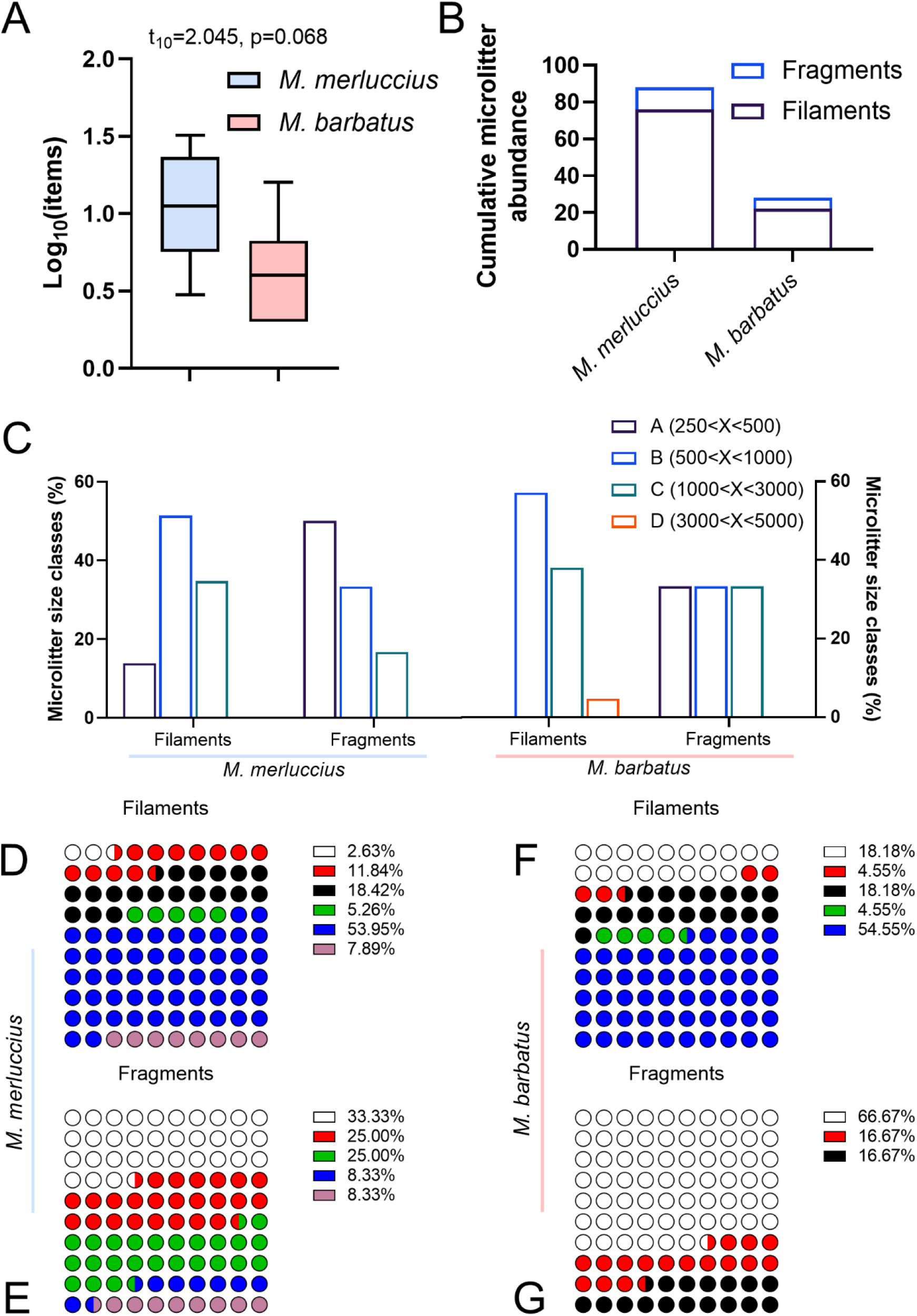
Quali-quantitative characterization of microlitter retrieved from fish stomach contents - A) Microlitter particle abundance distribution per species (log y scale). Whiskers plotted according to the Tukey method. B) Cumulative particle abundance per type. C) Size class distribution per microlitter shape type per species. D-G) Percentage of color abundance per microlitter shape type per species.

The retrieved microlitter was classified as filaments and fragments, and no films were found. 12.67 ± 4.27 filaments and 2 ± 0.71 fragments were retrieved from hakes, while 4.33 ± 2 filaments and 1.17 ± 0.37 fragments were found in mullets. Filaments were the most abundant shape type, with 76/88 particles (83.36%) in hake and 22/28 (78.57%) in mullet (Fig. 5B). B) *M. merluccius* ingested mostly filaments and fragments in the B and A size classes, respectively. The most represented filament size class in *M. barbatus* stomach contents was B, while fragments equally fell into the three size classes (Fig. 5C).

Filaments found in the stomach contents of *M. merluccius* were generally blue (53.95%) and black (18.42%) followed by red, green, white and other colored types (all below 12%). Blue, black and green filaments were found in similar percentages also in *M. barbatus* (54.55%, 18.18% and 4.55%, respectively). Microlitter fragments in hake and mullet were mostly white (33% and 66.67%, respectively) and red (25% and 16.67%, respectively); however, green, blue and other colored-fragments were retrieved only from hake, while black fragments were only found in mullet (Fig. 5D-G). Differences in microlitter stomach content by color between species was not statistically significant either for filaments (X^2^_(5)_ =9.38, p=0.094) or fragments (X^2^_(5)_ =5.63, p=0.34).

Forty items visually classified as plastics, equaling approximately the 35% of all items retrieved, were further analyzed by μFT-IR for a qualitative term of reference. Samples for which a spectrum could be obtained matched exclusively to synthetic polymers: filaments exclusively matched with HDPE while fragments were identified as HDPE, polyamide and polypropylene. HDPE and polypropylene, with a cumulative identification rate of 88.8%, were the two most frequent polymers. Selected spectra obtained from a blue fragment from *M. barbatus*, a blue filament from *M. merluccius* and a white fragment from *M. barbatus* are presented, showing a Pearson’s correlation coefficient of 0.87, 0.72 and 0.58 with reference spectra, respectively (Fig. 6 A-C).

**Fig. 6.**
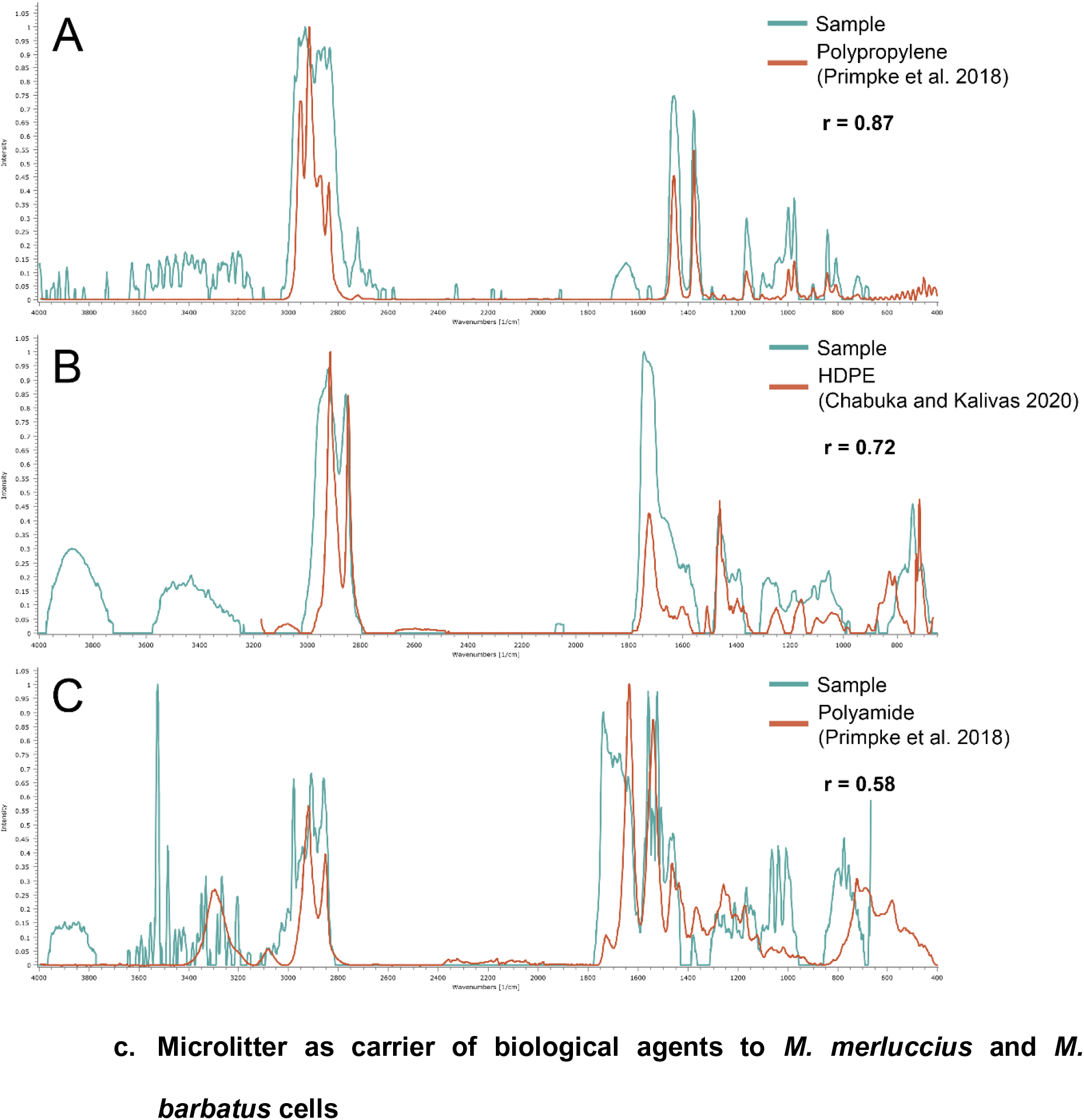
μFT-IR spectra - Spectra of randomly selected representative microlitter items retrieved from fish stomach contents. A) Blue fragment from *M. barbatus*; B) blue filament from *M. merluccius*; C) white fragment from *M. barbatus*. Matching with reference polymer as per the Open Specy open source database. r: Pearson’s correlation coefficient as measure of linear correlation between data sets.

### c. Microlitter as carrier of biological agents to *M. merluccius* and *M. barbatus* cells

To evaluate the presence of microorganisms and consequently validate the role of microlitter as carrier of biological agents, 10 μL cell suspension from all cultures following a 72-hour incubation were qualitatively assessed by optical microscopy. Bacilliform bacteria (grey arrowheads), unicellular fungi (white arrowheads) and flagellates (black arrowheads) were observed in microlitter-exposed cells (Fig. 7 B-D) but not in control cells (Fig. 7 A).

**Fig. 7.**
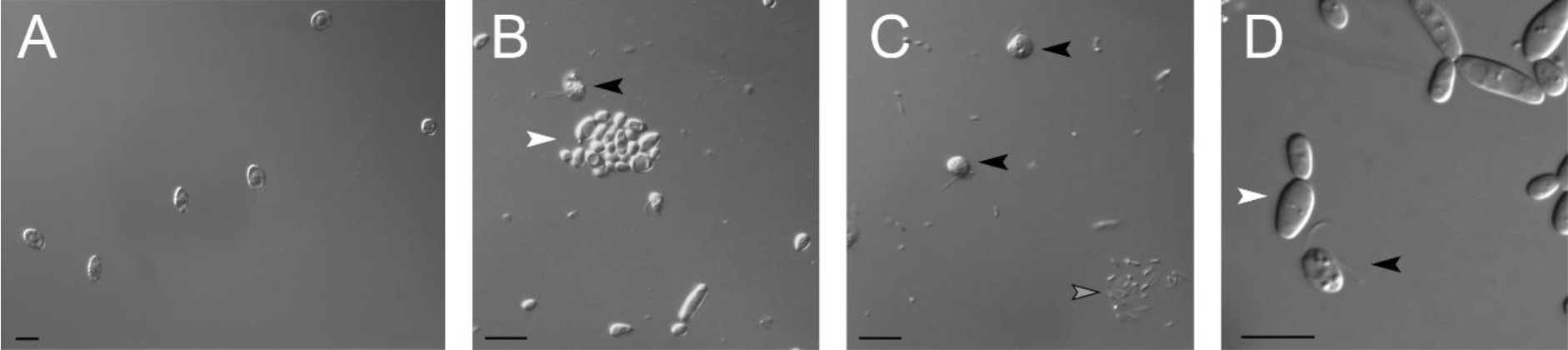
A-D Microlitter as a vector of biological agents - Examples of microorganisms observed in cell suspensions following a 72-hour incubation with microlitter. A) European hake splenic cell, negative control. B-D) Grey, white and black arrowheads indicate bacilliform bacteria, unicellular fungi and flagellates, respectively. Scale bars: 10 μm.

Their abundance was quantified over a 37.000 µm^2^ area and normalized to 100.000 µm^2^ area per species, organ and microlitter concentration. A variable degree of biological contamination was found in conditioned primary cultures of both species (Table 2).

For the hake, a significant effect of microlitter concentration on microbial counts was found only in HK primary cultures (H statistics=6.16, p=0.025). For the mullet, statistical significance was evidenced for microbial counts in G and HK (H statistics=7.62, p=0.036).

### d. Microlitter cytotoxicity in *M. merluccius* and *M. barbatus* cells

Microlitter cytotoxicity was evaluated based on the viability of cells from G, HK and SPL following a 2- and a 72-hour long incubation with two concentrations of microlitter sampled by 250 µm net tows (Fig. 8).

**Fig. 8.**
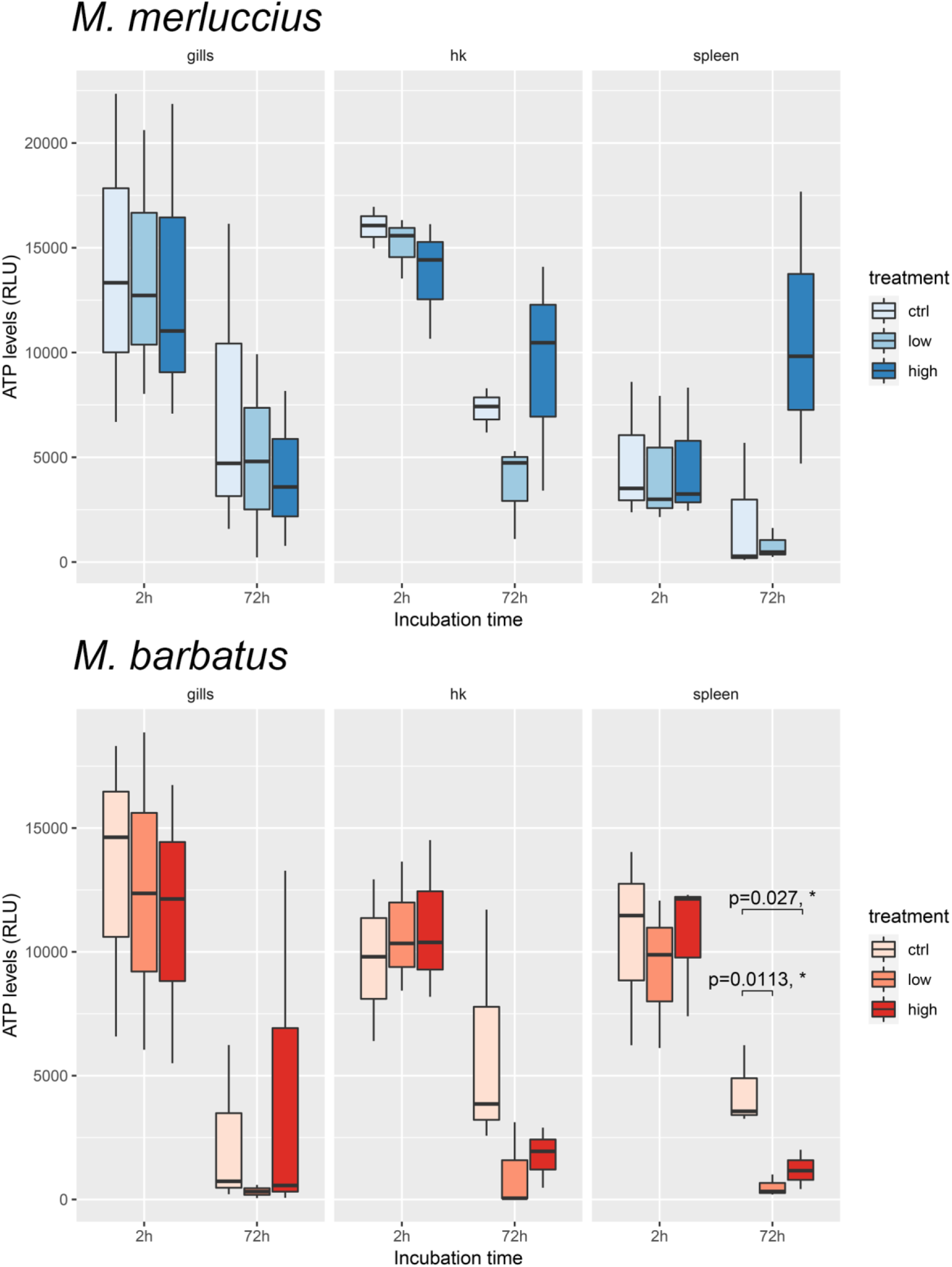
Quantification of intracellular ATP content as a proxy of cell viability - ATP data distribution per species, time, organ and treatment. Statistical significance as per one-way ANOVA followed by Tukey’s HSD *post hoc* test. *: *p*< 0.05

Exposure to microlitter did not induce any statistically significant decrease in intracellular ATP following the short incubation time in either species or organs (Table S3a-b).

In *M. merluccius*, microlitter induced a decrease in median cell viability after a 72-hour incubation in primary G cultures at the high concentration and in both HK and SPL cultures at the low concentration. The linear model fit to splenic cell culture data following the 72-hour exposure explained the 62.3% of ATP levels variation, even though differences among experimental groups were slightly non-significant (F(2,6) = 4.96, p=0.054) (Table S3a).

In *M. barbatus* primary cultures from all organs, the median intracellular ATP levels were lower in the Low and High groups than in corresponding controls. Such a decrease revealed a statistically significant main effect of microlitter concentration on splenic cells (F(2,6) = 10.8, p =0.01), with the overall treatment effect explaining almost 80% of ATP levels variation of the model (η^2^ = 0.783) (Table S3b). Pairwise comparisons between the control and the “Low” and “High” groups were statistically significant (p adjusted = 0.0113 and 0.027, respectively) (Fig. 8).

## 4. Discussion

In this work we applied a multidisciplinary approach combining oceanographical, spectroscopical, cellular and microscopical methods to characterize the quality and quantity of microlitter particles in the coastal surface waters and in the digestive tract of two commercially-valuable Mediterranean fish species; we also preliminarily addressed the cytotoxic potential of field-collected microlitter on primary cultures of cells extracted from mucosal (i.e. gills) and lymphoid (i.e. head kidney and spleen) organs.

The marine litter causes multiple environmental, economic, social, political and cultural impacts (Barboza et al., 2019; Galgani et al., 2019; GESAMP, 2015; UNEP, 2014), especially to the health and functioning of organisms and ecosystems (Corinaldesi et al., 2021; Garcia-Vazquez et al., 2018; Rios et al., 2007). At the European level, such pollutant was included among the 11 qualitative descriptors of the Marine Strategy Framework Directive against which the quality of the marine environment is assessed (European Parliament, 2008/56/EC). Extensive research has demonstrated the ubiquity of plastic pollution in several matrices such as beaches (Fortibuoni et al., 2021; Prevenios et al., 2018), sediments (Piazzolla et al., 2020; Renzi et al., 2018) and seawater (Atwood et al., 2019; Capriotti et al., 2021) – although remote (Cincinelli et al., 2017; Lusher et al., 2015). Microlitter was retrieved from all water samples taken within the framework of the PISCES project in a much higher (∼20-fold) average concentration (0.30 ± 0.02 items m^-^³) than litter particles > 5 mm (0.014 ± 0.003 items m^-^³). Keeping in mind the environmental and biological severity of litter < 5 mm, our results are in good agreement with microlitter concentrations reported from other areas of the Mediterranean Sea, Yellow Sea and oceanic waters (Baini et al., 2018; Cincinelli et al., 2017; Collignon et al., 2012; Constant et al., 2018; Cózar et al., 2015; de Lucia et al., 2018, 2014; Expósito et al., 2021; Fagiano et al., 2022; Fossi et al., 2012, 2016; Kazour et al., 2019; Lusher et al., 2015; Panti et al., 2015; Suaria et al., 2016; van der Hal et al., 2017; Zhang et al., 2017; Zhao et al., 2014) (Fig. 9), suggesting that also surveys that are not extensive in either duration or sample sizes can effectively capture the extent of microlitter pollution. This is desirable to minimize the impacts of research-related anthropogenic activities. An exception was represented by the Eastern Mediterranean Sea, which appears to be much more polluted than the western basin. We must highlight that data dispersion could not be quantified from Cózar et al. (2015) and Constant et al. (2018) as only mean items m^-3^ were reported, and that data from Vasilopoulou et al. (2021) was discarded because of non-informative results (41.31 ± 112.05 mean ± SD items m^-3^ - SD could be back-calculated from standard error because a sample size was clearly indicated by authors).

**Fig. 9.**
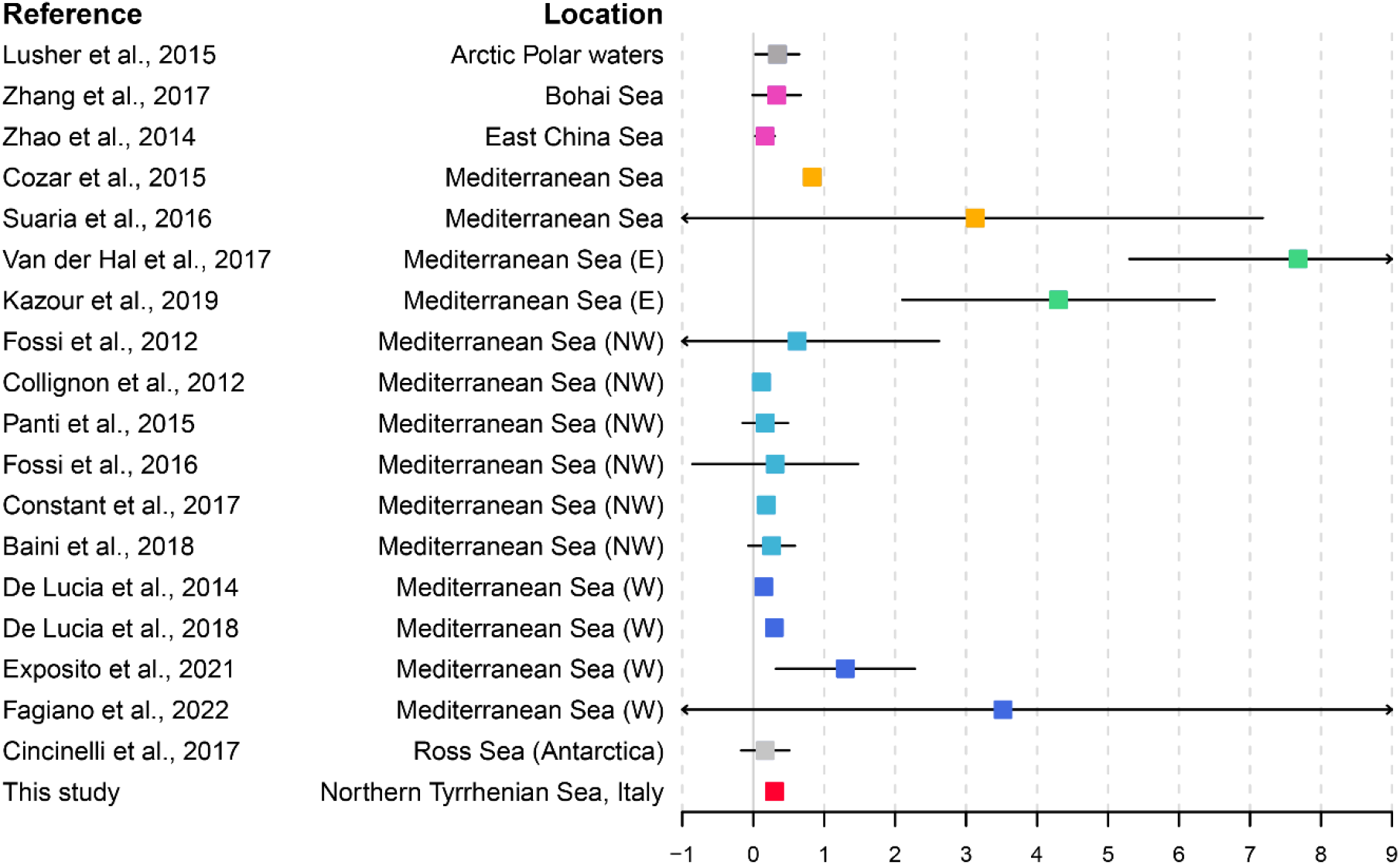
Comparison of microlitter densities retrieved from representative scientific literature with data herein presented. Boxes indicate mean ± SD items m^-3^. Sampling locations differentiated by color-coding.

The relationship between the marine biota and microlitter was so far mostly evaluated by ingestion rates (e.g. Rios-Fuster et al., 2019; Savoca et al., 2019). MPs are thought to be mistaken for or even purposefully chosen instead of food (Clark et al., 2016; Ling et al., 2017) probably also depending on their color (Du et al., 2021; Wright et al., 2013). The presence of both microlitter shape types in the stomachs of *M. merluccius* and *M. barbatus* and the lack of statistically significant differences based on the chromatic factor support the idea that microlitter may be ingested non-selectively by these two species, even though a biomagnification origin cannot be excluded. Hake and mullet were chosen as experimental models for a variety of reasons: on one hand they are among the most targeted demersal fish species by the Mediterranean deep-sea fisheries and the two most fished target species in the shallow area of the coastal sector (Sabatella et al., 2017; Tiralongo et al., 2021); on the other, they are regarded as bioindicators of coastal marine ecosystems and display a benthic feeding behavior at a certain life stage (juvenile *M. merluccius*) or throughout the lifespan (*M. barbatus*) (Carrozzi et al., 2019; Esposito et al., 2014). Moreover, some authors already described the occurrence of MPs in these two species (Atamanalp et al., 2021; Avio et al., 2020, 2015; Bellas et al., 2016; Digka et al., 2018; Giani et al., 2019; Mancuso et al., 2019). Here we confirmed that microlitter ranging in size from 250 to 3000 μm was present in the stomachs of both our target species in higher numbers compared to literature data (e.g. Avio et al., 2020; Bellas et al., 2016; Giani et al., 2019), especially for *M. Merluccius*. A reason for this may be found in an ontogenic trait of the species, for which a shift from benthic towards pelagic and necto-benthic feeding habits was demonstrated to occur over a TL of 18 cm, and in the fact that a greater microlitter contamination is usually observed in sediments than water column. Benthic macrolitter in the area was recently described quali-quantitatively, with the most abundant categories being ascribed to plastic (Mancini et al., 2021); MPs were also demonstrated to abound in superficial sediments in the study area (Piazzolla et al., 2020). The most commonly ingested microlitter items were confirmed by μFT-IR as high density polyethylene, polyamide and polypropylene (Chabuka and Kalivas, 2020; Primpke et al., 2018). On this, we must specify that a small number of spectra could be identified with certainty due to poor signal quality caused by the saturation of the detector (i.e. large sample size), or significant degradation of the plastic items likely caused by stomach acids. More importantly, we call the attention on the fact that such synthetic polymers were also demonstrated in marine sediments (Piazzolla et al., 2020) and in the atmosphere (Lucci et al., 2021) of the same area, pointing to the high and pervasive dispersion of anthropogenic litter across multiple environmental compartments.

It is known that MPs act as a carrier of biological agents (Amaral-Zettler, 2019; Kiessling et al., 2015), and our data confirmed this (Fig. 7). Because a dedicated experiment aimed at molecular taxonomy could not be set up due to limited availability of microorganisms on microlitter, flagellates were classified on phenotypic properties. Based on flagellar features and because they are extremely common in marine plankton, where they can be found free-swimming or attached to bacterial mats or other surfaces, we hypothesize they may belong to the *Paraphysomonas* or *Spumella* genus or to the aloricate Bicosoecida order. Microbial composition has the ability to condition the fate of MPs in the water column and sediments (Rogers et al., 2020). Once the microlitter is ingested, its associated microorganisms may colonize the gastrointestinal tract of the host, possibly affecting its welfare: in fact, harmful microorganisms, including potential human and animal pathogens, were found associated to litter (Zettler et al., 2013) and, according to data from Zwollo et al. (2021), serious consequences may arise due to the reduced ability to respond adequately to pathogens because of suboptimal humoral immune responses.

Research aimed at also elucidating physiological impacts of MPs have exposed fish to pristine commercially-available particles under controlled laboratory conditions. Their bioavailability was demonstrated and effects such as altered feeding behavior, metabolic disorders, energy depletion, growth impairment, delayed development, compromised immune response, reproduction and lifespan were reported (Botterell et al., 2019; Espinosa et al., 2019, 2017; Guerrera et al., 2021; Mazurais et al., 2015; Rios-Fuster et al., 2021; Sendra et al., 2021; Yong et al., 2020).

Recently, beach-sampled microlitter was employed in *in vivo* experiments on the European sea bass *Dicentrachus labrax* (Zitouni et al., 2021) and medaka *Oryzias latipes* (Pannetier et al., 2020) to investigate survival, development, uptake, oxidative stress and genotoxicity following the administration of a microlitter-spiked feed. Their results showed the ability of environmental MPs to i) accumulate in fish organs, ii) significantly affect the activity of enzymes involved in the antioxidant defense system and iii) induce DNA damages following acute exposures. HK primary cultures were also employed to explore the impacts of non-environmental MPs on the abundance and antibody response of B cells in rainbow trout (Zwollo et al., 2021): a lower rate of B cell development together with reduced expression of Ig heavy chain genes were found, suggesting that not only innate but also adaptive immunity may be threatened by such an emerging contaminant.

Despite some similarities with the three just-mentioned studies may be perceived, we must highlight that no other research has ever investigated the apical cytotoxicity event in primary cell cultures derived from fish mucosal and lymphoid organs following their exposure to microlitter collected in the same area from where animals were sampled (literature search conducted on Web of Science on December 3, 2021). The primary organs for examining the cytotoxic effects of microlitter would have been the stomach or intestine, but the methods for cell extraction are difficult, lengthy, prone to contamination and poor in yield, which were all incompatible factors with our experimental design. Keeping in mind that microlitter and MP impacts occur via cell internalization or chemical contamination and that the former mode of action was not expectable due to the large size of particles employed, we investigated additional immunologically-relevant organs that could be reached by the release of chemical contaminants contained by or adsorbed on the microlitter particles, and affected in their physiological status. We believe that our results, obtained in an attempt to bridge the fields of biological oceanography and experimental toxicology, are biologically significant and actual because i) microlitter particles and fish specimens originated from the same sampling site, ii) microlitter cytotoxicity was measured by the well-established, highly-sensitive and unambiguous direct luciferase-based quantification of cellular ATP (Cree and Andreotti, 1997; Mahto et al., 2010) iii) primary cultures were obtained from organs that are key in ensuring immune barrier and competency and iv) the suitability of the strategy for testing for MP toxicity was overall demonstrated and recently reviewed in details (Revel et al., 2021). In addition, fish have been increasingly established as experimental models in the fields of biomedical sciences and toxicology because they share many similarities with higher vertebrates immunology-wise (Miccoli et al., 2021a; Scapigliati et al., 2018).

Taking into account cytotoxicity data (Fig. 8) and the lack of statistically different microbial counts observed within *M. barbatus* spleen cultures (Table 2), splenic cell subpopulations appeared to be the most sensitive to microlitter exposures among all investigated organs. No further reduction in ATP levels were seen in the High compared to the Low condition, suggesting that such a pollutant can impact cell viability already at concentrations that are in line with estimates modelled over the next three decades. These results are concerning because spleen, together with thymus and kidney, is the major lymphoid organ of teleosts in which adaptive immune responses are mounted (Flajnik, 2018; Zapata et al., 2006). Neither the physiological endpoints reported in the large majority of scientific literature nor our results herein presented provide insight into the molecular mechanisms underlying microlitter toxicity pathways, but inform about apical events manifested either by the whole organism or primary cell cultures, respectively. However, the novelty of our approach was to provide data on a lower, possibly more predictive, level of biological organization (cellular vs. organismal) by means of so-called New Approach Methodologies, which heavily rely on *in vitro* testing. This is compliant with the 3Rs principle in animal testing, in addition to having been validated by the latest internationally-agreed test guidelines (OECD, 2021) and supported by regulatory toxicology roadmaps (e.g. [EPA’s strategic vision](https://www.epa.gov/pesticide-science-and-assessing-pesticide-risks/strategic-vision-adopting-new-approach)).

## 5. Conclusions

In conclusion, the present study has investigated the anthropogenic litter in the coastal epipelagic Northern Tyrrhenian Sea and the stomach of two commercially-relevant fish species, validated the microlitter fraction as a carrier of biological agents and, for the first time, demonstrated that splenic cell viability is negatively affected following exposure to such a contaminant. Future investigations with larger sample sizes, primary or continuous cell cultures from additional organs and more in-depth methodological approaches are warranted for clarifying the susceptibility of *Merluccius merluccius* and *Mullus barbatus* to anthropogenic microlitter.

## 6. Author contribution

AM: Conceptualization, Funding acquisition, Data curation, Formal analysis, Visualization, Supervision, Project administration, Writing - original draft, Writing - review.

EM: Conceptualization, Funding acquisition, Investigation, Writing - Review & Editing.

PRS: Methodology, Investigation, Writing - original draft, Writing - Review & Editing.

GDV: Methodology, Resources, Writing - Review & Editing.

GS: Resources, Supervision, Writing - Review & Editing.

SP: Supervision, Writing - Review & Editing

## 7. Acknowledgements

This research was funded by the ‘‘PISCES’’ project (FEAMP PO 2014–2020 promoted by Lazio Region, Italy) and the ‘‘Departments of Excellence-2018’’ Program (Dipartimenti di Eccellenza) of the Italian Ministry of Education, University and Research awarded to the DIBAF Department of University of Tuscia. AM and EM are extremely thankful to the ‘‘Marinai & Caratisti’’ Cooperative of Civitavecchia and to Dr. Roberto Arciprete for their support in all sampling operations. Financial support to GDV was provided by the Grant to Department of Science, Roma Tre University (MIUR-Italy Dipartimenti di Eccellenza, ARTICOLO 1, COMMI 314-337 LEGGE 232/2016). Dr. Daniele Piazzolla of the Laboratory of Experimental Oceanology and Marine Ecology (University of Tuscia) is acknowledged for his contribution to the PISCES project. AM wishes to thank Prof. Pierangelo Luporini (University of Camerino) for his valuable input on flagellate taxonomy.

---

Table S1 Sea surface microlitter abundance and density (items m^-3^) per color per type per net tow and relative share of litter types.

Table S2 Sea surface litter size classes per particle type per net tow.

Table S3 Full statistical details of the one way ANOVA tests performed on cytotoxic data of C) *Merluccius merluccius* and B) *Mullus barbatus*, per time and organ. DFn: degrees of freedom in the numerator; DFd: degrees of freedom in the denominator; F: test statistic for ANOVA; ges: generalized eta squared.

