## Supplemental Table 2 for "First evidence of *in vitro* cytotoxic effects of marine microlitter on *Merluccius merluccius* and *Mullus barbatus*, two Mediterranean commercial fish species"

| | SUPR1 | | | | | SUPR2 | | | | | SUPR3 | | | | | SUPR4 | | | | |
|---|---|---|---|---|---|---|---|---|---|---|---|---|---|---|---|---|---|---|---|---|
| | A (250<X<500) | B (500<X<1000) | C (1000<X<3000) | D (3000<X<5000) | E (X>5000) | A (250<X<500) | B (500<X<1000) | C (1000<X<3000) | D (3000<X<5000) | E (X>5000) | A (250<X<500) | B (500<X<1000) | C (1000<X<3000) | D (3000<X<5000) | E (X>5000) | A (250<X<500) | B (500<X<1000) | C (1000<X<3000) | D (3000<X<5000) | E (X>5000) |
| **Filaments** | 3 | 10 | 14 | 6 | 3 | 4 | 4 | 22 | 3 | 0 | 0 | 0 | 3 | 6 | 4 | 0 | 11 | 30 | 8 | 4 |
| **Fragments** | 6 | 5 | 3 | 1 | 0 | 1 | 9 | 9 | 8 | 2 | 9 | 30 | 20 | 8 | 1 | 1 | 4 | 9 | 0 | 1 |
| **Films** | 9 | 26 | 21 | 4 | 0 | 3 | 8 | 13 | 4 | 2 | 6 | 11 | 24 | 8 | 0 | 5 | 38 | 24 | 5 | 0 |
