## Supplemental Table 3a for "First evidence of *in vitro* cytotoxic effects of marine microlitter on *Merluccius merluccius* and *Mullus barbatus*, two Mediterranean commercial fish species"

| Time | Organ | Effect | DFn | DFd | F | *p* | *p*<0.05 | ges |
|---|---|---|---|---|---|---|---|---|
| **2h** | gills | treatment | 2 | 6 | 0.009 | 0.991 | | 0.003 |
| **2h** | hk | treatment | 2 | 6 | 1.076 | 0.399 | | 0.264 |
| **2h** | spleen | treatment | 2 | 6 | 0.017 | 0.983 | | 0.006 |
| **72h** | gills | treatment | 2 | 6 | 0.278 | 0.767 | | 0.085 |
| **72h** | hk | treatment | 2 | 6 | 2.031 | 0.212 | | 0.404 |
| **72h** | spleen | treatment | 2 | 6 | 4.96 | 0.054 | | 0.623 |
