## Supplemental Table 3b for "First evidence of *in vitro* cytotoxic effects of marine microlitter on *Merluccius merluccius* and *Mullus barbatus*, two Mediterranean commercial fish species"

| Time | Organ | Effect | DFn | DFd | F | *p* | *p*<0.05 | ges |
|---|---|---|---|---|---|---|---|---|
| **2h** | gills | treatment | 2 | 6 | 0.061 | 0.941 | | 0.02 |
| **2h** | hk | treatment | 2 | 6 | 0.16 | 0.856 | | 0.051 |
| **2h** | spleen | treatment | 2 | 6 | 0.141 | 0.871 | | 0.045 |
| **72h** | gills | treatment | 2 | 6 | 0.623 | 0.568 | | 0.172 |
| **72h** | hk | treatment | 2 | 6 | 2.244 | 0.187 | | 0.428 |
| **72h** | spleen | treatment | 2 | 6 | 10.805 | 0.01 | * | 0.783 |
